# Voxelwise encoding models of body stimuli reveal a representational gradient from low-level visual features to postural features in extrastriate body area

**DOI:** 10.1101/2022.12.19.521151

**Authors:** Giuseppe Marrazzo, Federico De Martino, Agustin Lage-Castellanos, Maarten J. Vaessen, Beatrice de Gelder

**Affiliations:** Department of Cognitive Neuroscience, Faculty of Psychology and Neuroscience, Maastricht University, Limburg 6200 MD, Maastricht, The Netherlands; Department of Computer Science, University College London, London WC1E 6BT, UK; Center for Magnetic Resonance Research, Department of Radiology, University of Minnesota, Minneapolis, MN 55455, United States and Department of NeuroInformatics; Cuban Center for Neuroscience, Street 190 e/25 and 27 Cubanacán Playa Havana, CP 11600, Cuba

## Abstract

Previous research has focused on the role of the extrastriate body area (EBA) in category-specific body representation, but the specific features that are represented in this area are not well understood. This study used ultra-high field fMRI and banded ridge regression to investigate the coding of body images by comparing the performance of three encoding models in predicting brain activity in ventral visual cortex and specifically the EBA. Our results suggest that EBA represents body stimuli based on a combination of low-level visual features and postural features.

**Author Summary:** Historically, research on body representation in the brain has focused on category-specific representation, using fMRI to investigate the most posterior body selective region, the extrastriate body area (EBA). However, the role of this area in body perception is still not well understood. This study aims to clarify the role of EBA, in coding information about body images. Using ultra-high field neuroimaging (fMRI) and advanced encoding techniques we tested different computational hypotheses to understand how body images are represented in EBA. Our results suggest that EBA represents bodies using a combination of low-level properties and postural information extracted from the stimulus.

## Introduction

Faces and bodies are amongst the most frequently encountered visual objects and provide essential information about the behaviour of conspecifics. In contrast to face perception, body perception is still poorly understood. Mainstream research on body representation in humans has focussed on category specific body representation in the brain, investigated with fMRI to identify conceptual category defined functional selectivity. Initially a body category selective area was reported in the middle occipital\temporal gyrus, the extrastriate body area (EBA) (1). Later a second body selective area was described in the fusiform cortex and labelled fusiform body area (FBA) (2). Studies on body representation in nonhuman primates using fMRI as well as invasive electrophysiology resulted in a similar situation of multiple body sensitive patches in temporal cortex (3). Once multiple category selective areas were reported in human as well as in nonhuman primate, the central issue is to understand how body images are coded in the different body selective areas and how to account for the observed body selectivity.

An attractive notion that has been explored but ultimately not supported is that EBA coded body parts and the more anterior FBA whole bodies, but this distinction proved inconclusive (for review, (4, 5)). An earlier proposal that EBA was selective for body parts and the more anterior FBA for whole bodies and their overall configuration (6, 7) is not supported by current findings in humans or non-human primates (3, 5). Furthermore, this is not easy to combine with findings that activity in EBA is influenced by task setting (8-10) but also by experimental manipulations of semantic attributes like gender and emotional expression (11-18). The fact that such stimulus attributes have an impact on the level of activity observed in EBA also challenges the notion that EBA only codes for body parts.

Thus our current understanding of how body images are processed shows a gap between the extraction of low-level physical features of the stimulus taking place in early visual cortex and the generation of a high-order semantic concept of bodies at stake in processing information about emotions or action and presumably linked to FBA activity (5). In view of its location in temporal cortex it is likely that the kind of coding to expect in EBA is related to computing some subsymbolic body features rather than identifiable body parts because the latter already implies high level body category representations (19). Candidate subsymbolic features are overall shape representation and related to that, viewpoint tolerance, an important dimension in the posterior to anterior gradient of object recognition. Studies in non-human primates that use single cell recordings indicate that moving from posterior to anterior temporal cortex, body patch neurons increase their selectivity for body identity and posture, while there is a decrease in viewpoint selectivity. Specifically, recordings in body selective patches, middle superior temporal body (MSB) and anterior superior temporal body (ASB) showed strong viewpoint selectivity for the former and conversely, high tolerance for the latter (20). Furthermore, Caspari and colleagues using the same set of category stimuli as Kumar and colleagues, showed similar decoding pattern between monkeys and humans in body selective regions, suggesting an homology between the human EBA and monkey MSB as well as the human FBA and monkey ASB (20, 21).

This suggests a general principle of object coding in the inferior temporal cortex (IT): a greater tolerance to image transformations that preserve identity (22) and, in the case of bodies, posture, for more anterior patches. The monkey data fits human fMRI work that found viewpoint-invariant decoding of body identity in FBA but not EBA (14), but as noted above, results of between-area differences in fMRI multi voxel pattern analysis (MVPA) are difficult to interpret (23).

An important question is whether a similar posterior to anterior organisation can be found for EBA by using ultra high field fMRI in combination with computational hypotheses. One popular approach to test and compare different computational hypotheses of brain function is to use (linearized) encoding (24, 25) approaches. In these approaches brain activity (e.g. the blood oxygen level dependent (BOLD) signals in a voxel or an brain area in fMRI) is predicted from the features of (different) computational models, and their accuracy can be compared to adjudicate between competing models or partitioned with the respect to the variance explained by each of the models (26-32).

We used ultra-high field fMRI and linearized encoding to evaluate to what extent the response in extrastriate body areas can be explained on the one hand by low-level visual features (Gabor) (33) and on the other by the features extracted by two computational models that represent the postural features of the body (kp2d, kp3d) (34, 35) (see Material and Methods).

## Results

### Behavioural analysis

The analysis of the responses to the questionnaires revealed that no action was recognised for 92% of the stimuli (298 out of 324). Likewise, no emotion was recognised for 97% of the stimuli (314 out of 324). Participants reported that they focused on the overall body pose in 65% of the cases (211 out of 324), on the hands in 20% (64 out of 324) of the cases and on the arms for 11% (38 out of 324). The full report on the behavioural results is found in the supplementary material.

### Univariate analysis and voxels selection for encoding

In each subject, voxels that significantly (q(FDR)<0.05) responded to the localizer conditions (main effect) were selected for the encoding analysis (Fig. 2a). At the group level, we observed significant (q(FDR<0.01) activation in occipital-temporal cortex as well as parietal cortex in the occipital gyrus (superior/middle/inferior) (SOG/MOG/IOG), fusiform gyrus (FG), lingual gyrus (LG), middle temporal gyrus (MTG), superior parietal lobule (SPL), intraparietal sulcus (IPS), inferior temporal sulcus (ITS), lateral occipital sulcus (LOS), superior temporal sulcus (STS) (Fig 2b).

**Figure 1.**
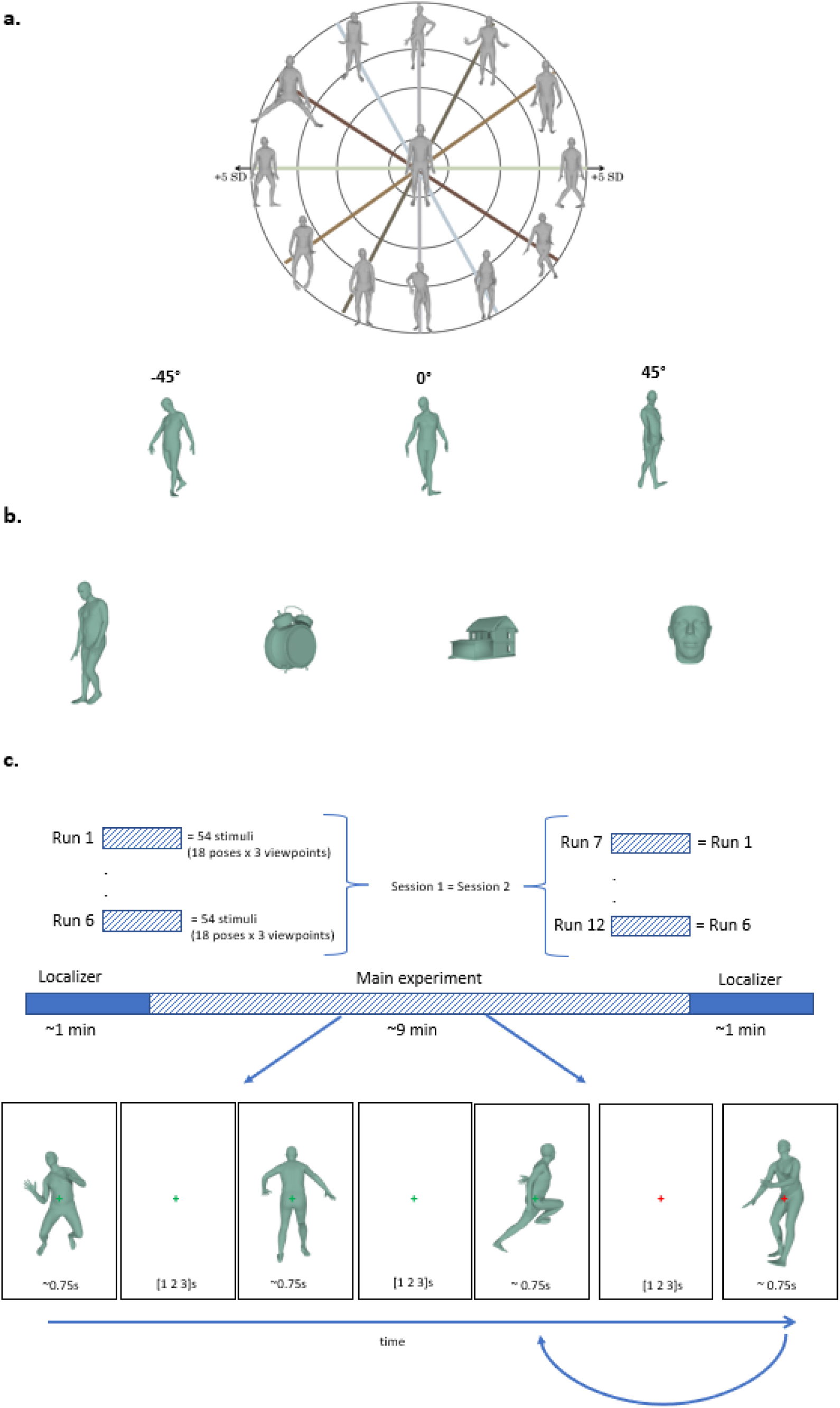
Stimuli and experimental procedure. **(a)** (top) Stimuli were generated by randomly sampling the latent space of the VAE (34, 35). The 32-dimensional latent space was sampled in three shells, defined by the value of the standard deviation from the mean pose, to ensure variability among the generated poses. (a) (bottom) 108 unique poses were generated from three different viewpoints: 0º (frontal), -45º (left rotated) and 45º (right rotated) for a total of 324 unique stimuli. **(b)** Body sensitive areas were identified by mean of a localizer using stimuli selected from four different object categories: bodies, tools, houses, and faces. These stimuli underwent the same rendering process as the stimuli of the main experiment. **(c)** During the main experiment participants performed a one-back task. They fixated on the green cross and were presented with pictures of body poses each for approximately 750 ms followed by a blank screen which appeared for 1, 2 or 3 s. When the fixation cross turned red, they had to report by button press whether the current stimulus matched the previously presented one.

**Figure 2.**
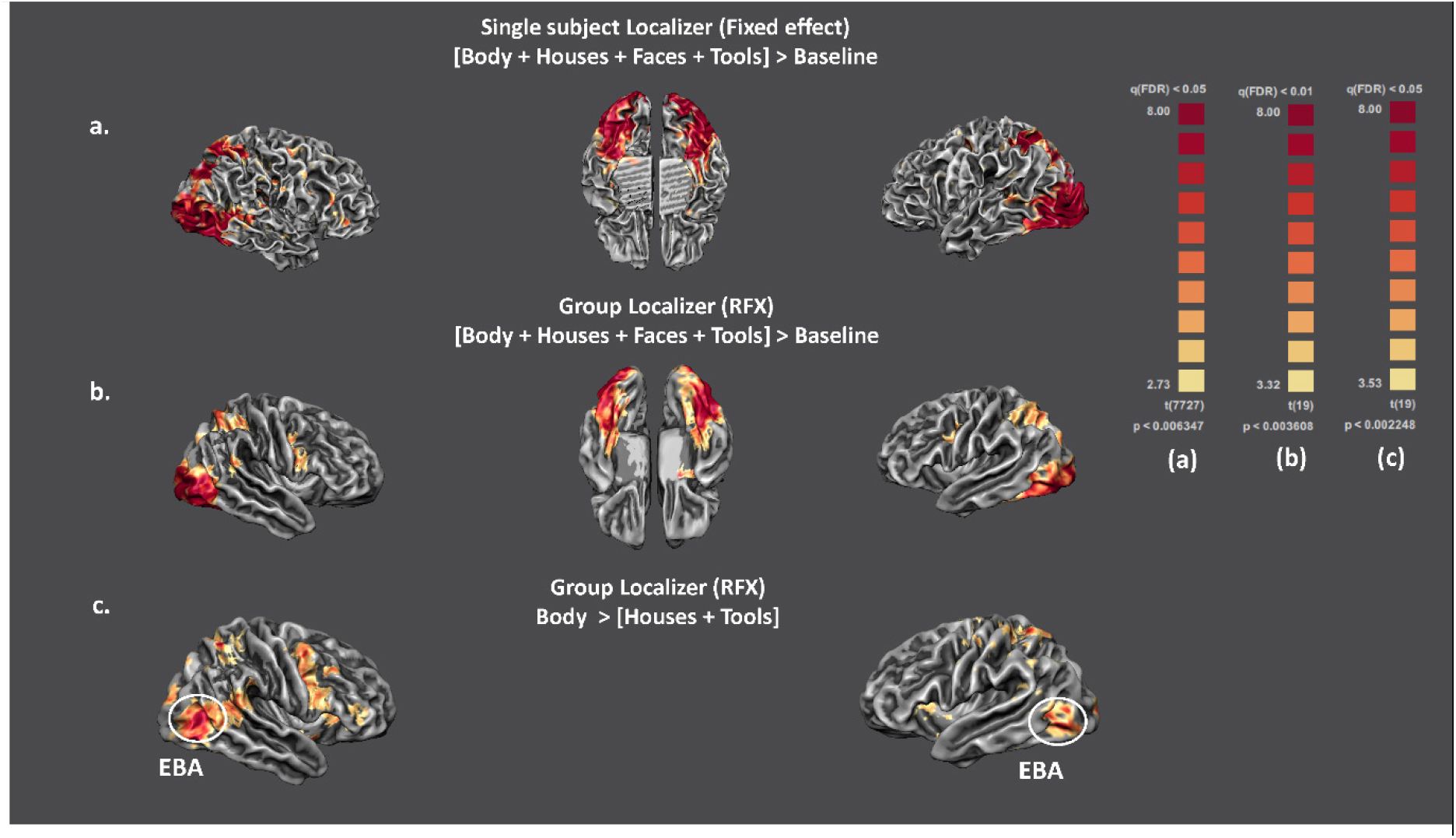
Univariate analysis. **(a)** Brain maps showing the responses for the main effect of the localizer in a single subject computed with a fixed-effect GLM. This map was created in volume space (q(FDR)<0.05) and overlaid on the subject mesh for visualization purposes. **(b)** Brain activation for the main effect of the localizer obtained when including all the subjects in a RFX GLM. The activation map is corrected for multiple comparison at q(FDR)<0.01 and is cluster thresholded (cluster size = 25). **(c)** Body selective regions obtained by contrasting the localizer conditions Body > Objects (Houses + Tools). As in (a), the statistical thresholding of the map was performed in volume space (q(FDR)<0.05) and then overlaid on the group average mesh for visualization purposes. We used this contrast to obtain a group definition of EBA which was intersected with single subjects’ activation maps for the subsequent ROI analysis.

Subtracting the response to object stimuli from the response to body stimuli allowed us to define EBA. This cluster spanned the MOG, MTG as well as the ITS (Fig 2c). The voxels selection for the encoding analysis was performed at the individual level and based on the main effect. A probabilistic map (computed by counting the number of subjects for which a given voxel was included in the analysis) showed a consistent overlap with the functionally defined EBA (Fig 3).

**Figure 3.**
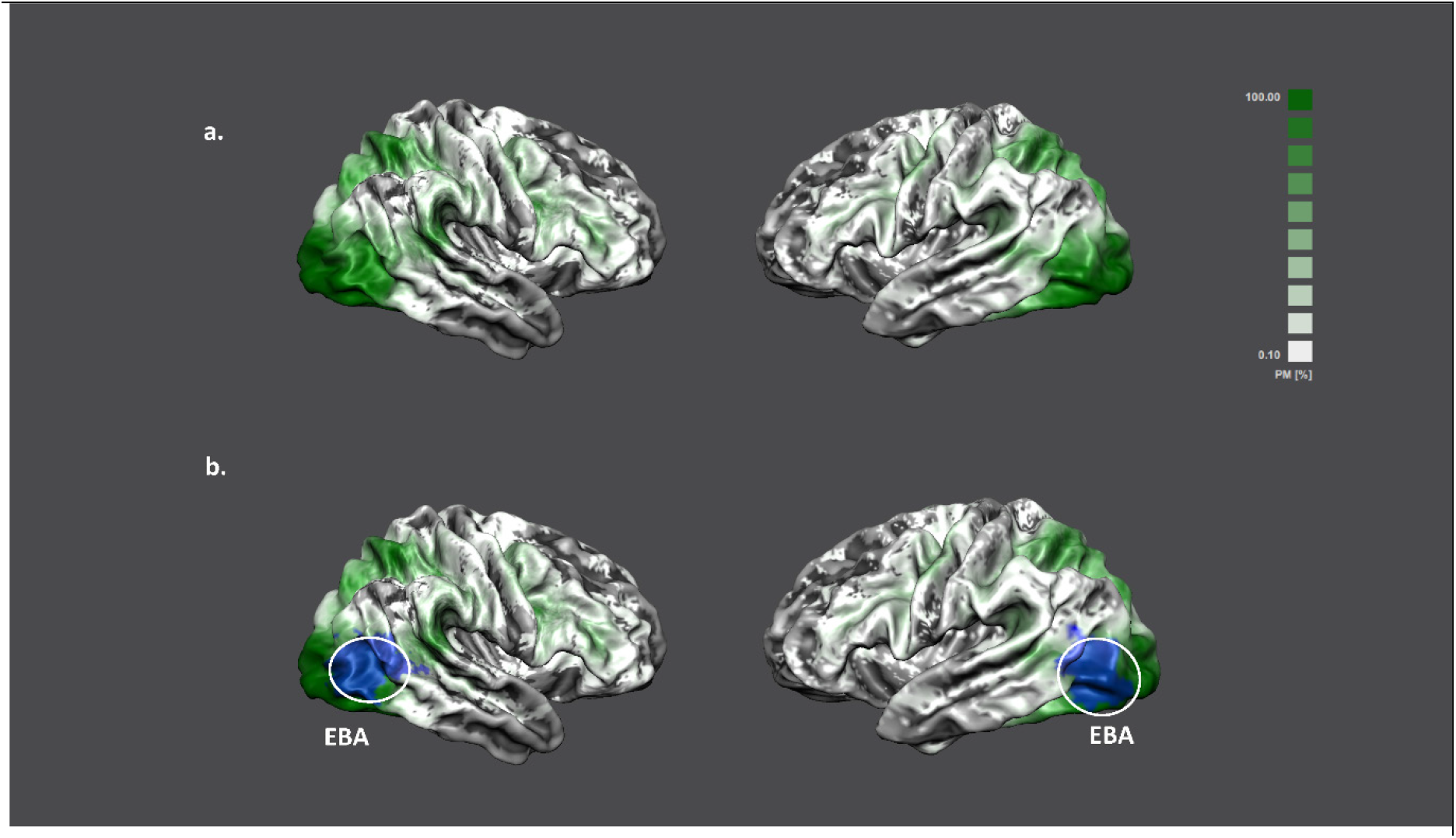
Probabilistic map of the main effect of the localizer. **(a)** This brain map shows the extent of the overlap between participants within the main effect of the localizer computed for each participant across all runs via a fixed-effect GLM. This overlap is expressed via a probability map where at each spatial location the percentage of the relative number of subjects leading to significant activity is reported (low probability → high probability: white → green). **(b)** In the second row, we overlay the binarized (suprathreshold voxels q(FDR)<0.05 = 1) group definition of EBA (in blue) (see Fig. 2c) on the probabilistic map. This shows that most (90-100%) of the participants shared significant responses (q(FDR)<0.05) within the region of interest.

### Encoding results

The voxels selected using the response to the localizer were submitted to the encoding analysis. The response to the body stimuli presented in the main experiment (data independent from the localizer) were modelled using banded ridge regression. The group performance of the joint (three) encoding model is shown in Fig. 4. The accuracy of the joint (kp2d, kp3d, Gabor) encoding model at the group level (after statistical testing and correction for multiple comparisons) is shown in Fig. 4. We found that when combining information from the three models we could significantly predict responses to novel stimuli (Fig. 4) throughout the ventral visual cortex (SOG, MOG, IOG, ITG, MTG, FG, LOS), and in parietal cortex (SPL). We assessed spatial differences in how models contributed to the fMRI response by colour coding the relative contribution of each of the models to the overall prediction accuracy (Fig. 5). The response to bodies in early visual cortical areas was in average better explained by the Gabor model (blue-purple-dark magenta).

**Figure 4.**
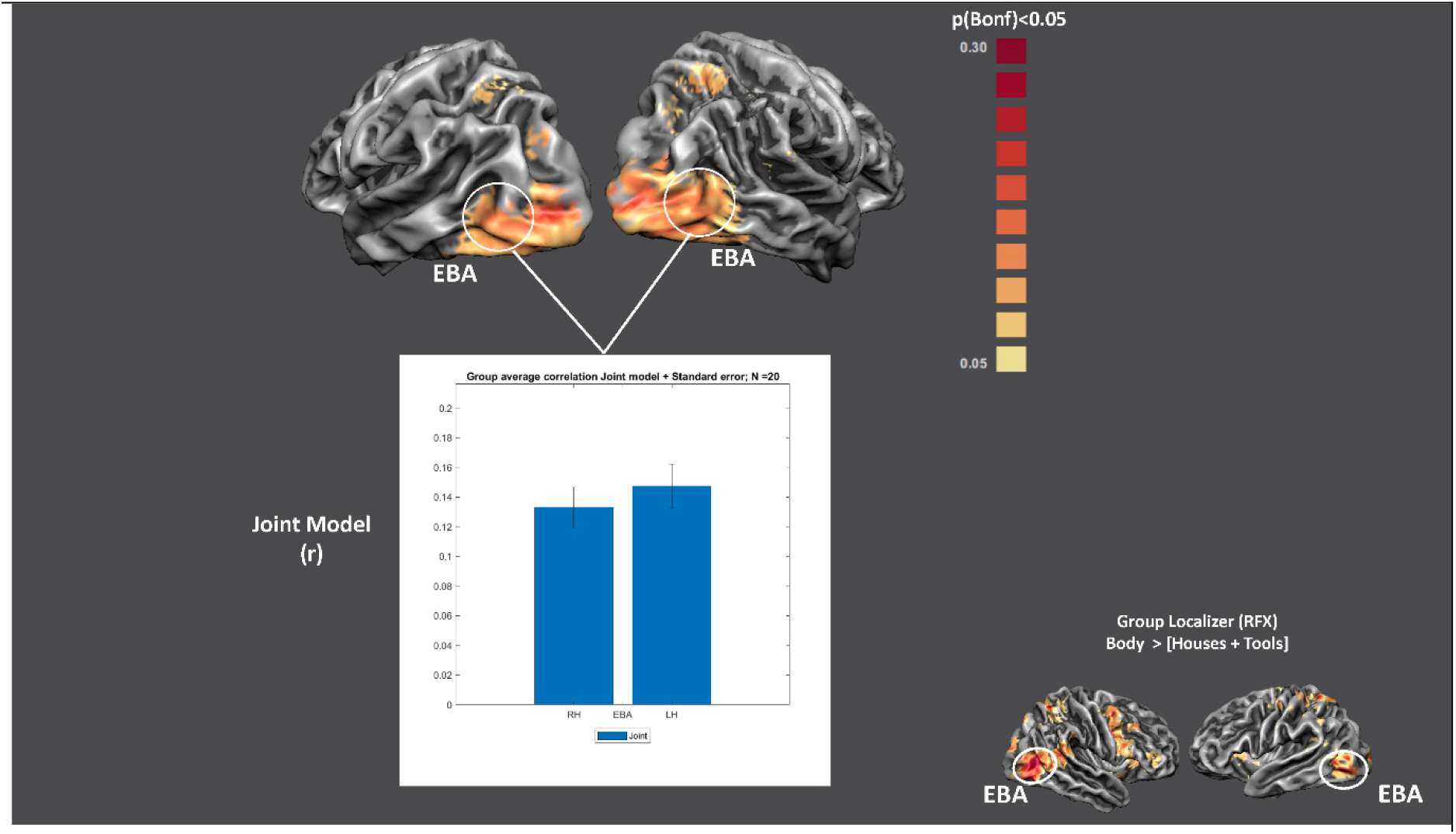
Joint model performance. Group Prediction accuracy for the joint model (kp2d, kp3d, Gabor). Statistical significance was assessed via permutation test (subject wise sign-flipping, 10000 times), and correction for multiple comparison was performed using Bonferroni correction (p<0.05). The bar plot depicts the group (mean + standard error) correlation coefficient between the joint model predictions and brain response to novel stimuli (test stimuli) across participants in bilateral EBA. We did not find any significant difference across hemispheres (two-sample t-test, p=0.481). For reference, the bottom right corner shows the functional definition of EBA already presented in Fig. 2c.

**Figure 5.**
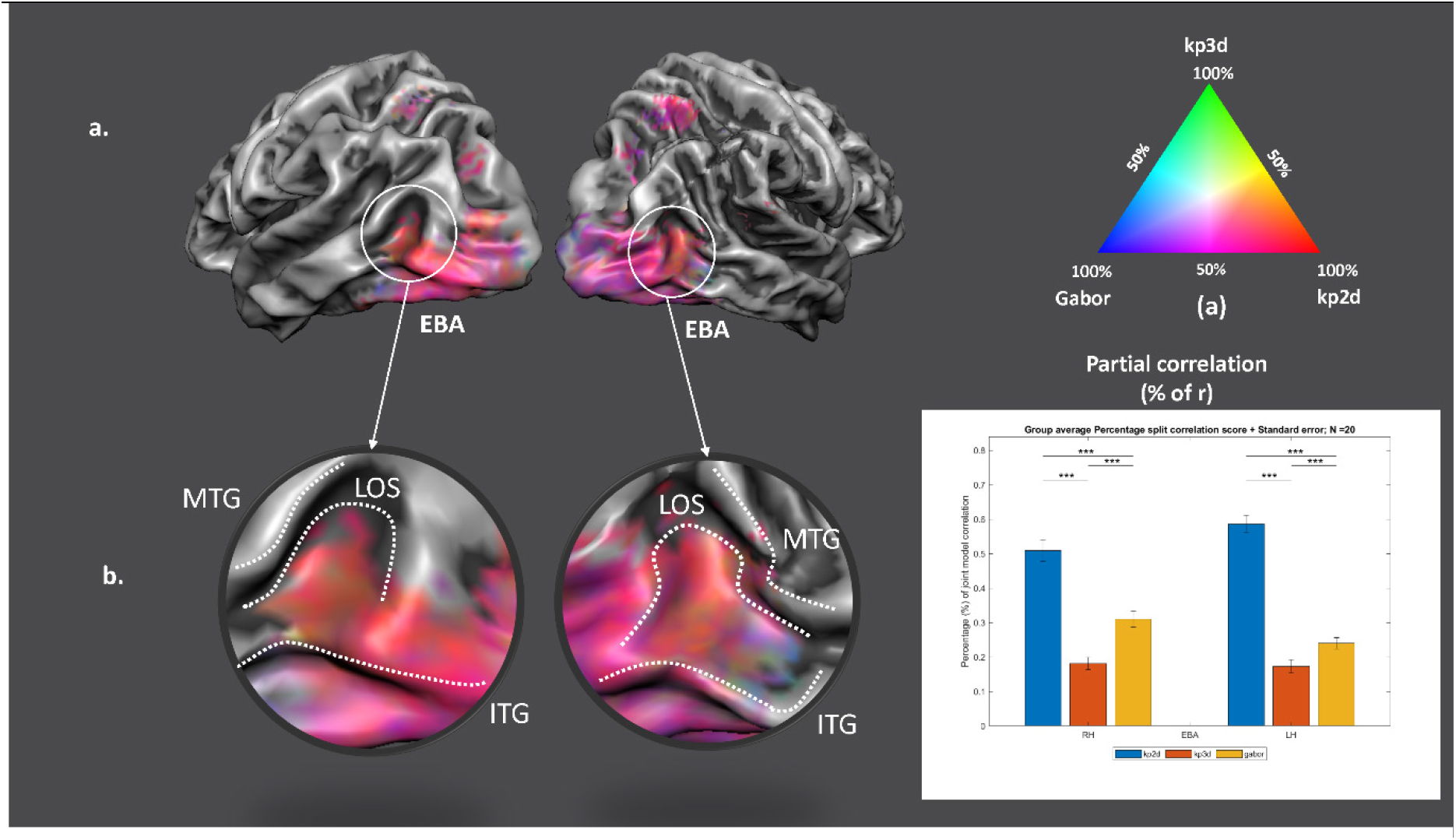
Comparison between encoding models. **(a)** RGB map in which each vertex is colour coded according to the relative contribution of each model to the accuracy of the joint model (red = 100% kp2d; blue = 100% Gabor; green = 100% kp3d). **(b)** In EBA, the information contained in the joint model predictions which significantly correlates with BOLD activity is split across models with kp2d accounting for 50-60% of the variance, Gabor approximately 25-30% of the variance and kp3d the remaining 15-20%. We tested for statistical difference across models’ pair (solid lines at the top), using a two-sample t-test (*** p<0.0001) (see bar plot). Additionally, the variance explained follows a gradient from the posterior part (posterior ITG/LOS) to the anterior (anterior LOS) of EBA, with darker shades of magenta in the posterior part indicating higher representation of low-level body features (Gabor), and lighter shades of magenta in the anterior part indicating higher representation of mid-level features (kp2d-kp3d). ITG = inferior temporal gyrus; MTG = middle temporal gyrus; LOS = lateral occipital sulcus.

Moving to higher visual cortical areas corresponded to a shift in the relative contribution towards a combination of kp2d and Gabor (magenta), while in EBA the model that contributed most to the prediction accuracy was kp2d (magenta - light magenta - pink). When considering EBA (Fig. 5), the joint model significantly predicted brain responses to test stimuli (Fig. 4), and the kp2d model accounts for approximately 50-60% of the variance of this prediction (Fig. 5b). It is worth noting that when considering the spatial distribution of relative model contributions to the prediction accuracy (Fig. 5), the posterior part of EBA, specifically the posterior part of lateral occipital sulcus (LOS) was best explained by the Gabor model (dark magenta area), while the anterior part of LOS showed lighter shades of magenta indicating that the leading representation is kp2d.

## Discussion

In this study, we used ultra-high field fMRI to determine the main (stimulus) features that drive brain responses to still body stimuli, with a particular focus to the responses in the extrastriate body selective area (EBA). We compared the performance of three encoding models using banded ridge regression. We observed that a combination of the three models (kp2d, kp3d, Gabor) could significantly predict fMRI BOLD responses in ventral cortex and in parietal cortex (SPL). The partial correlation analysis revealed that, in EBA, approximately 50% of the variance of the prediction accuracy is explained by kp2d, 30% by Gabor and 20% by kp3d. These results lead us to conclude that EBA represents body stimuli based on the combination of low-level visual features and postural features.

EBA was originally defined as a category selective area associated with body representation but the computations underlying this selective response are not yet well understood. Previous proposals stressed the role of EBA for individual body parts but not whole body images (1, 36, 37). These results are difficult to combine with evidence that EBA is selective for human bodies when only represented as stick figures, line drawings or silhouettes (38). Our findings are consistent with the latter hypothesis as the kp2d/3d model explain approximately 70% of the accuracy in EBA.

The Gabor model proposed by (33) was specifically constructed to encode low-level visual features such as spatial frequency, location, size and object orientation. Gabor based models have been shown to be powerful tool for inferring (encoding/decoding) (25, 26, 28, 30, 31, 33, 39-42) brain activity inside and outside early visual cortex. Our findings suggest that the variance explained by the Gabor model shows a decreasing gradient from early to higher-level visual cortex. This suggests that within early sensory regions (superior occipital gyrus, blue patches in Fig. 4b) Gabor features are critical for predicting BOLD responses to body stimuli. Conversely, the variance explained by kp2d shows the opposite gradient, and it is highest in EBA. This suggests that postural features are critical in driving the response to body pictures in EBA. Interestingly, the transition between low-level features driving the response in early areas and mid-level (postural) features driving the response in high-level visual cortex (EBA) at the group level is smooth and suggests a dynamic, stimulus dependent, representation of bodies (5). Likewise, similar patterns can be seen at the single subject level (see Supplementary material).

Another important point is the performance difference between kp2d and kp3d. These models represent body poses as the spatial location of specific keypoints (joints, hand, head etc). In the case of kp3d, the keypoints represent the 3D coordinates used by VPoser to pose the mesh (34, 35) and construct the actual stimulus. Similarly, kp2d represents the orthogonal projection of the 3D coordinates on the camera plane. Therefore, the only difference between the models is that kp3d is isotropic (invariant across viewpoints), whereas the features of kp2d change across different view of the same pose. Our findings show that between kp2d/3d, banded ridge almost always selects the former as predictive and consider the latter as redundant. This is reflected in the percentage maps where on average kp2d outperforms kp3d. One possible explanation for this result is that the information contained in the 3^rd^ dimension of kp3d was not needed to explain the variance in the data and, as a result, the selected feature space was kp2d for most of the voxels, suggesting that the viewpoint information is encoded in EBA. Previous research has shown that EBA is sensitive to body orientation (11-14, 16), although we did not find significant differences in brain activity when looking at differences between viewpoints (RFXGLM with three viewpoints as predictors of interest). This result is in line with what has been shown in single cell recordings on primates, where the MSB (analogous of the EBA in humans) showed strong viewpoint selectivity (20).

It is worth mentioning that our stimuli were specifically controlled for the presence of high-level stimulus attributes (i.e. emotion, action information) and validated using behavioural ratings (see behavioural analysis). Many previous studies have shown that activity in EBA is modulated by emotional body expressions (43-50). Moreover, a recent study has shown that unique information from the posture feature limb contractions is involved in fearful body expression perception (51). This indicates that body expression may be based on body posture and movement features rather than implicating body representation as a high-level semantic category (5).

Our results corroborate the notion that the functional EBA definition spans several anatomical regions with potentially different roles. Specifically, the EBA may be subdivided in three anatomical regions (52) located respectively in the inferior temporal gyrus (ITG), middle temporal gyrus (MTG) and lateral occipital sulcus (LOS). When looking closely at the partial correlation patterns in EBA around the anatomical landmarks ITG, MTG, LOS (Fig. 5b and the barplot Fig. 5b) we see that not all the variance can be explained by combining the kp2d model with the Gabor model. This is graphically represented in figure 5b, where we find a green (or green derived) colour in the anterior part of EBA (anterior LOS/ITG), indicating that the variance explained by kp3d model is on average located in the more anterior portion of EBA. Specifically, bodies in the anterior portion of EBA are represented as a combination of kp2d/Gabor features with the kp3d model (yellow/light-blue patches in Fig. 5b). This finding might indicate that, as shown for early sensory regions, body representation in EBA is differentially encoded, going from a low-level representation (Gabor like/blue patches) in pITG/pLOS, to a mid-level (viewpoint dependent) postural representation (kp2d, light-magenta, orange, pink patches) in the (middle) LOS to a high-level (viewpoint independent) postural representation (kp3d) in aITG/aLOS (green, light-blue, yellow patches).

Concerning the other major body selective region FBA, we observed that for the voxels significantly responding to localizer stimuli, the group definition of this region was not consistent across participants. Moreover, among the voxels functionally identified as part of the FBA, only few survived the statistical correction for multiple comparison of the encoding analysis. For completeness, we include the results of the encoding compared to EBA in the supplementary information. Briefly, the joint model performs significantly worse in FBA than in EBA, this could be due to low signal to noise ratio in the area. Nonetheless, the barplot depicting the percentage of the correlation explained by each model reveals a similar behaviour to what has been presented for EBA. The main difference is that kp3d model has an increase (from 20 to 25%) in percentage of correlation explained in FBA, at the expense of the correlation explained by kp2d. This is consistent with the fact that FBA has higher viewpoint tolerance than EBA as is expected if FBA is more involved in higher cognitive processing of body information like personal identity (13-15).

Taken together, these results suggest that the EBA encodes features pertaining specifically to posture. This representation appears to be viewpoint dependent posteriorly (pITG/pLOS) whereas greater viewpoint tolerance arises anteriorly (aITG/aLOS). On this account, the body selectivity observed in many studies in EBA is rooted in body specific feature representation that is not yet dependent of high order body categorisation processes. Future research must investigate whether these body selective features are rooted in uniquely defined biomechanical constraints, in human skeleton keypoint priors or also in sensorimotor processes.

## Material and methods

### Participants

20 right-handed subjects (8 males, mean age = 24.4 ± 3.4 years) participated in this study. They all had normal (or corrected to normal) vision and were recruited from Maastricht University student cohorts. All subjects were naïve to the task and the stimuli and received monetary compensation for their participation (7.5 € VVV vouchers/per hour or a bank transfer for the same amount; 4h in total, 30 €). Scanning sessions took place at the neuroimaging facility Scannexus at Maastricht University. All experimental procedure conformed to the Declaration of Helsinki and the study was approved by the Ethics Committee of Maastricht University.

### Stimuli

#### Main experiment stimuli

The stimulus set consisted of 108 pictures of 3D rendered body meshes shown in different orientations: 0º (frontal), -45º (left rotated) and 45º (right rotated) for a total of 324 unique images. Examples of the stimuli in the different orientation are shown in Fig 1a. 3D rendered body meshes were generated via VPoser, a variational autoencoder (VAE) trained to learn a 32-dimensional (normal distribution) latent representation of Skinned Multi-Person Linear Model (SMPL) parameters (34, 35). The stimuli used in the study were generated via randomly sampling the latent space and generating via the decoder part of the VPoser the associated body image. To also sample images sufficiently distant from the mean image (and thus maintain a sufficiently large variability of poses in the stimulus set), we sampled the latent space within three distinct shells defined by the standard deviations from the mean pose (Fig. 1a). Ultimately, the body images were generated by transferring the decoded SMPL parameters to a posed mesh. The resulting body poses had mean widths and heights of 2.43º x 5.22º of visual angle and were colour rendered (mean RGB: 120,157,144).

#### Localizer stimuli

Stimuli for the localizer experiment consisted of 3D rendered images depicting four object categories: faces, bodies, tools, and houses (Fig. 1b). The stimuli were colour rendered using the same colour for the main experiment stimuli (mean RGB: 120,157,144). None of the stimuli from the localizer were used in the main experiment.

### Behavioural validation

Stimuli used in the main experiment were generated via the VAE. Random sampling from the latent space allowed us to produce a varied set of body poses but did not allow us to control the stimuli for the possible presence of semantic body attributes like action or emotion. Therefore, we asked 113 participants (25 excluded for missing data, 88 in total: 29 males, mean age = 23 ± 4 years, 72 right handed) to rate the stimuli using a questionnaire consisting of both categorical and likert-scale questions. Participants were presented with 1/3 (108) of the total stimuli (324) for 750 ms each. For each participant, the stimuli were pseudo-randomized (108 stimuli randomly selected for each participant, but evenly distributed so that each stimulus got approximately the same number of answers). After each presentation, participants were asked to answer 6 questions about the emotional expression, action content; salience of specific body parts; implied body movement and realism of the posture (see supplementary material).

### MRI acquisition and experimental procedure

Participants viewed the stimuli while lying supine in the scanner. Stimuli were presented on a screen positioned behind participant’s head at the end of the scanner bore (distance screen/eye = 99 cm) which the participants could see via a mirror attached to the head coil. The screen had a resolution of 1920×1200 pixels, and its angular size was 16º (horizontal) x 10º (vertical). The experiment was coded in Matlab (v2018b The MathWorks Inc., Natick, MA, USA) using the Psychophysics Toolbox extensions (53-55).

Each participant underwent two MRI sessions, we collected a total of twelve functional runs (six runs per session) and one set of anatomical images. Images were acquired in a 7T MR scanner (Siemens Magnetom) using a 32-channel (NOVA) head coil. Anatomical images were collected via a T1-weighted MP2RAGE: 0.7 mm isotropic, repetition time (TR) = 5000 ms, echo time (TE) = 2.47 ms, matrix size= 320 × 320, number of slices = 240. The functional dataset covered the entire brain and was acquired via T2*-weighted Multi-Band accelerated 2D-EPI BOLD sequence, multiband acceleration factor = 3, voxel size = 1.6 mm isotropic, TR = 1000 ms, TE = 20 ms, number of slices = 68 without gaps; matrix size = 128 × 128.

Each run consisted of three main sections: 1) two short localizers parts (approximately one minute at the beginning and at the end of the run), during which images were presented in blocks of categories (faces, bodies, tools, and houses), and 2) a main experimental part where stimuli (body images different from the ones used in the localizer) were presented following a fast event-related design. Participants were asked to fixate at all times on the green cross at the centre of the screen.

Each localizer block each contained six images which were presented for 750 ms and followed by 250 ms blank screen. Each block lasted six seconds followed by a fixation period of eight seconds and each category block was presented once at the beginning and once at the end of each run (24 blocks per condition across the 12 runs). During the localizer participants did not perform any task.

During the main experiment, stimuli were presented for 750 ms with an inter stimulus interval that was pseudo-randomised to be 1, 2 or 3 TRs. To keep attention on the stimuli, participants performed a one-back task om stimulus identity. Following a visual cue (colour change of the fixation cross), they were asked report via a button press whether the current stimulus was the same as the previous one (Fig. 1c). Within each run, the experimental section consisted of the presentation of 54 stimuli (18 unique poses x 3 viewpoints) repeated 3 times each. Six target trials were added for a total of 168 trials. Across the two sessions each of the 108 unique poses were repeated 18 times (3 repetitions x 3 viewpoints x 2 sessions) across the 12 runs, whereas the 324 unique stimuli were repeated 6 times (3 repetitions x 2 sessions).

Preprocessing was performed using BrainVoyager software (v22.2, Brain Innovation B.V., Maastricht, the Netherlands) and FSL (56-58). The following steps were performed in BrainVoyager unless indicated otherwise. EPI Distortion was corrected using the Correction based on Opposite Phase Encoding (COPE) plugin in BrainVoyager, where the amount of distortion is estimated based on volumes acquired with opposite phase-encoding (PE) with respect to the PE direction of the main experiment volumes (59), after which subsequent corrections is applied to the functional volumes. Other preprocessing steps included: scan slice time correction using cubic spline, 3D motion correction using trilinear/sinc interpolation and high-pass filtering (GLM Fourier) cut off 3 cycles per run. During the 3D motion correction all the runs were aligned to the first volume of the first run. Anatomical images were resampled at 0.5mm isotropic resolution using sinc interpolation and then normalized to Talairach space (60). To ensure a correct functional-anatomical and functional-functional alignment, the first volume of the first run was coregistered to the anatomical data in native space using boundary based registration (61). Volume Time Courses (VTCs) were created for each run in the normalized space (sinc interpolation) and exported in nifti format for further processing in FSL. To further reduce non-linear intersession distortions, functional images were additionally corrected using the fnirt command in FSL (62) using as template the first volume of the first run in normalized space. Prior to the encoding analysis (and following an initial general linear model [GLM] analysis aimed at identifying regions of interest based on the response to the localizer blocks), we performed an additional denoising step of the functional time series by regressing out the stimulus onset (convolved with a canonical hemodynamic response function [HRF]) and the motion parameters.

Segmentation of white matter (WM) and gray matter (GM) boundary was performed in BrainVoyager using the deep learning-based segmentation algorithm and in house Matlab scripts. The resulting boundaries were then inflated to a reference sphere and aligned using cortex based alignment (CBA) (63). The aligned meshes were averaged to create a group WM-GM mesh for each hemisphere.

### Voxels selection for encoding analysis

The functional time series of each participant were analysed using a fixed-effect GLM with 5 predictors (4 for the localizer blocks and 1 modelling the responses to all the stimuli in the main experiment). Motion parameters were included in the design matrix as nuisance regressors. The estimated regressor coefficients representing the response to the localizer blocks were used for voxel selection. A voxel was selected for the encoding analysis if significantly active (q(FDR)<0.05) within the main effect of the localizer (Body, Houses, Tools, Faces – Fig. 2). Note that this selection is unbiased to the response to the main stimuli presented in the experimental section of each run.

To assess the spatial consistency of activation to the localizer across subjects, we created a probabilistic functional map depicting, at each spatial location, the percentage of subjects for which that location was significantly (q(FDR)<0.05) modulated by the localizer blocks (Fig 3a).

### Functional ROI definition

We defined body selective regions at the group level using a random-effect GLM (RFX-GLM), in which EBA was defined using the localizer contrast Body > Objects([Houses + Tools]) (64) with a statistical threshold of q(FDR) < 0.05. Functional images from every participant were spatially smoothed using a gaussian filter (FWHM = 4mm) and then entered the RFX GLM in which we defined 5 predictors of interest (4 for the localizer 1 for modelling the responses to the main experiment). For each participant, we regressed out signals coming from head motion by including motion parameters in the design matrix. Responses from each subject were selected via intersection with the group ROI definition of EBA and the single subject localizer’s main effect map (see previous paragraph). Figure 3b projects the group definition of EBA onto the probabilistic functional map of the localizer’s main effect.

The group level body sensitive ROIs were intersected with the single subject activation maps (see previous section) to obtain individual ROIs. Note again that while this procedure makes use of the same data (localizer) twice, its purpose was to define single subject regions to be subsequently used for encoding analysis which was performed on an independent portion of the data set. Figure 3b reports the overlap between EBA defined at the group level and the probabilistic activation maps of the localizer’s main effect.

### Encoding models

In order to understand what determines the response to body images we tested several hypotheses, represented by different computational models, using fMRI encoding (24, 25, 29, 65). We compared the performance (accuracy in predicting left out data) of three encoding models. The first represented body stimuli using the position of joints in two dimensions (kp2d) using 54 keypoints (joints, hand and facial features like eyeballs, neck and jaw) plus one keypoint for global rotation extracted during the stimulus creation using VPoser (35). This encoding model extracts for each pose the orthogonal projection of the pose’s spatial coordinates on the camera plane which ultimately constitutes the image coordinates (i.e. x,y) of the keypoints. Therefore, this model has 110 features (55 kp * 2 dimensions). The second model represented body stimuli using the three-dimensional position of the keypoints (kp3d) extracted from VPoser. This representation differs from the kp2d one by adding the third dimension (no projection on the camera plane), resulting in an encoding model with 165 features (55 kp * 3 dimensions). It is important to note that the main difference between the kp2d and kp3d representations is that the latter is viewpoint invariant as the position of the joints is independent from the angle under which the object is observed.

The last encoding model we tested is a Gabor filtering of the images (33, 66-68). In this procedure, each stimulus was transformed into the Commission internationale de l’éclairage (CIE) L*A*B* color space and the luminance signals then passed through a bank of 1425 spatial Gabor filters differing in position, orientation, and spatial frequency (33, 69, 70). Ultimately, the filters output underwent a logarithmic non-linear compression in which large values were scaled down more than small values. For details on this procedure we refer to the original publication (33).

### Banded ridge regression and model estimates’

Generally, in the linearized encoding framework (as applied in fMRI) the information explained in brain activity is obtained via L2-regularized (ridge) regression (71). Ridge regression is a powerful tool which allows to improve performance of encoding models whose features are nearly collinear, and it minimizes overfitting. When dealing with more than one encoding model, ridge regression can either estimate parameters of a joint feature space (combining all feature spaces in one encoding model) or obtain model estimates from each encoding model separately. Fitting a joint model with ridge regression allows considering the complementarity of different feature spaces but subjects all models (feature sets) to a unique regularization. As the optimal regularization required when fitting each individual feature space may differ (since it depends, among others, on factors such as number of features and features covariances) (27), fitting a joint model with one regularization parameter may be suboptimal and can be extended to banded ridge regression. In banded ridge regression, separate regularization per parameters for each feature space are optimized, which ultimately improves model performance by reducing spurious correlations and ignoring non-predictive feature spaces (27, 28). In the present work we used banded ridge regression to fit the three encoding models and performed a decomposition of the variance explained by each of the models following established procedures (27). All analyses were performed using a publicly available repository in Python (Himalaya, https://github.com/gallantlab/himalaya).

Model training and testing were performed in cross-validation (3-folds: training on 8 runs [216 stimuli] and testing on 4 runs [108 stimuli]). For each fold, the training data were additionally split in training set and validation set using split-half crossvalidation. Within the (split-half) training set a combination of random search and gradient descent (27) was used to choose the model (regularization strength and model parameters) that maximized the prediction accuracy on the validation set. Ultimately, the best model over the two (split-half) folds was selected to be tested on the yield out test data (4 runs). The fMRI predicted time courses were estimated as follows. Within each fold, the models’ representations of the training stimuli were normalized (each feature was standardized to zero mean and unit variance withing the training set). The feature matrices representing the stimuli were then combined with the information of the stimuli onset during the experimental runs. This resulted in an experimental design matrix (nrTRs x NrFeatures) in which each stimulus was described by its representation by each of the models. To account for the hemodynamic response, we delayed each feature of the experimental design matrix (15 delays spanning 15 seconds). The same procedure was applied to the test data, with the only difference that when standardizing the model matrices, the mean and standard deviation obtained from the training data were used. We used banded ridge regression to determine the relationship between the fMRI response at each voxel, which significantly responded to the localizer stimuli (p(FDR)<0.05), and the features of the encoding models (stimulus representations).

For each cross-validation, we assessed the accuracy of the model in predicting fMRI time series by computing the correlation between the predicted fMRI response to novel stimuli (4 runs, 108 stimuli) and the actual responses. The accuracy obtained across the three folds were then averaged. To obtain the contribution of each of the models to the overall accuracy we computed the partial correlation between the measured time series and the prediction obtained when considering each of the models individually (27).

### Group maps and statistical inference

To evaluate the statistical significance of the model fittings, accuracy maps of each subject were projected on the cortex based aligned group WM-GM mesh. We computed the probability of the mean accuracy (across subjects) to be higher than chance by sign flipping (10000 times) the correlations. This procedure allowed estimating a non-parametric null distribution for each vertex, which was used to obtain a significance value for the mean accuracy. We accounted for the multiple comparisons by correcting the p-values using Bonferroni correction (i.e. dividing by the number of tests, equal to the number of vertices in the analysis).

## Acknowledgements

The authors are grateful to Rufin Vogels and Giancarlo Valente for fruitful discussion and comments on the original draft. The authors declare no competing financial interests.

## Data availability statement

Data and code are being prepared to be shared.

## CRediT authorship contribution statement

**Giuseppe Marrazzo:** Conceptualization, Investigation, Software, Formal analysis, Validation, Visualization, Writing – Original Draft Preparation, Writing - review & editing. **Federico De Martino:** Conceptualization, Investigation, Supervision, Validation, Writing - review & editing. **Agustin Lage-Castellanos:** Conceptualization, Supervision, Validation, Writing - review & editing. **Maarten J. Vaessen:** Conceptualization, Investigation, Software. **Beatrice de Gelder:** Conceptualization, Project Administration, Resources, Supervision, Funding Acquisition, Writing – Original Draft Preparation, Writing - review & editing

